# CMLD-2 Attenuates Myofibroblast Activation and Bleomycin-Induced Pulmonary Fibrosis in Mice through Antagonizing ELAVL1-Mediated Osteopontin mRNA Stabilization

**DOI:** 10.1101/2022.03.20.484975

**Authors:** Guo Qiongya, Ren Hongyan, Li Meng, Liu Lifan, Li Wenting, Zhang Jingjing, Wang Xiaoli, Hu Yiping, Zang Kaixuan, An Yunxia, Li Lin, Wei Li, Xu Zhiwei, Guo Zhiping, Ulrich Costabel, Zhang Xiaoju, Wang Zheng

**Affiliations:** Department of Digestive Diseases, the People’s Hospital of Zhengzhou University; Department of Respiratory and Critical Care Medicine, the People’s Hospital of Zhengzhou University; Henan University People’s Hospital; Department of Tropical Medicine, the Second Affiliated Hosptial of Hainan Medical College; Department of Respiratory and Critical Care Medicine, Zhou Kou Central Hospital; Zhengzhou University People’s Hospital; Emergency Department, the People’s Hospital of Zhengzhou University; Department of Thoracic Surgery, the People’s Hospital of Zhengzhou University; Department of Clinical Research and Biostatistics, the People’s Hospital of Zhengzhou University; Department of Radiology, Fuwai Central China Cardiovascular Hospital; Department of Pneumonology and Rare Lung Diseases, Ruhrlandklinik, Duisberg-Essen University

**Keywords:** RNA binding protein, HuR/ELAVL1, Pulmonary fibrosis, Fibroblast, Myofibroblast, Fibroblast-myofibroblast transition (FMT), Transforming growth factor β (TGF-β), Osteopontin, CMLD-2

## Abstract

**Background:** Fibroblast-myofibroblasts transition (FMT) is one of the hallmark cellular processes in pulmonary fibrosis. This study is to investigate the effects of CMLD-2 in FMT and pulmonary fibrosis, which antagonizes HuR, a supposedly key regulatory RNA binding protein (RBP).

**Methods:** HuR or other deferentially expressed RBPs during TGF-β1-induced FMT were analyzed by transcriptomic methods, and further validated *in vitro* or in fibrotic lung specimens. The effects of HuR overexpression, down-regulation or inhibition by an antagonist CMLD-2 were analyzed in FMT or bleomycin-induced experimental lung fibrosis. HuR-targeting RNA and their interactions were analyzed by CLIP-seq.

**Results:** HuR, hnRNPA1, hnRNPE1, TIA1 and TFRC were significantly up-regulated, while ESRP1, ESRP2 and TTP were significantly down-regulated. Cytoplasmic expression of HuR was activated in IPF lung tissue and rat lungs of bleomycin-induced fibrosis. HuR overexpression induced α-SMA and collagen I expression, increased the proliferation and migration capacities of fibroblasts with or without the stimulation of TGF-β1. HuR knockdown by shRNA inhibited the proliferation, transition, collagen production and migration properties in fibroblasts or in TGF-β1-stimulated myofibroblasts. Combinative analysis of RNA-seq and CLIP-seq data revealed major HuR binding motifs and several HuR-bound, differentially expressed mRNAs in FMT, specifically SPP1 mRNA encoding osteopontin. HuR-mediated SPP1 mRNA stabilization was further validated by RIP-PCR and half-life analysis using SPP1 mutant transcripts. Inhibiting HuR using CMLD-2 attenuated SPP1/osteopontin expression, TGF-β1-induced FMT *in vitro* and bleomycin-induced pulmonary fibrosis in mice. Nuclear-cytoplasmic shuttle of HuR was activated in TGF-β1-induced FMT, which was inhibited by p38MAPK (SB203580) or PKC (Go-6976) inhibition, but not associated with phosphorylation of HuR.

**Conclusions:** Fibroblast-myofibroblast transition is activated by HuR-SPP1 mRNA interactions, and CMLD-2 is potentiated to be a therapeutic agent targeting HuR for fibroblast-myofibroblast transition and pulmonary fibrosis.

## 1. Introduction

Pulmonary fibrosis, the endpoint sequalae of interstitial lung diseases (ILD) or lung damage, has been emerging as a major cause of respiratory disability and death, and is becoming progressively prevalent worldwide ^1,2^. Treatment with pirfenidone or nintedanib has been shown to slow, if not prevent or reverse, on patients with idiopathic pulmonary fibrosis (IPF) or non-IPF fibrotic lung diseases ^2^. Except for pirfenidone and nintedanib, there are few recommended drugs proven to be effective in phase III clinical trials that enrolls patients with pulmonary fibrosis ^3,4^. In this scenario, novel preventative and therapeutic targets and approaches are desperately needed for patients with pulmonary fibrosis ^4^. Success will require improved understanding of fundamental mechanisms that control the pathogenesis of pulmonary fibrosis.

Current conceptual pathological process of fibrosis consists of aberrant injury, aging, inflammation, repair and eventually the replacement of normal lung parenchyma by dense fibrous scar ^5,6^. Epithelial cells, fibroblasts, macrophages, lipofibroblasts, endothelial cells, pericytes and immune cells all play important roles in the progressive process of fibrosis ^7^. As one of the most important effector cell type, fibroblasts are activated and producing excessive extracellular matrix ingredients ^8,9^. Myofibroblasts are abundantly accumulated in fibrotic lung tissue, especially fibroblast foci ^10^. As revealed by single-cell multi-omic studies, myofibroblasts originate and transit from resident interstitial fibroblasts, pericytes, macrophages or lipofibroblasts, which is named as fibroblast/pericyte/macrophage/lipofibroblast-myofibroblast transition. These myofibroblasts that has undergone cellular transition processes are characterized by α-SMA overexpression and gain of motility, rendering them an overactivated cell type that produces even more extracellular ingredients and profibrotic cytokines than fibroblasts ^9,11,12^. The process of fibroblast-myofibroblast transition (FMT) is mediated by multiple signals, among which TGF-β1 is most widely studied. Inactivation of TGF-β1 signaling inhibits fibroblast activation, FMT, and the process of pulmonary fibrosis ^13^. Elucidation of TGF-β1-induced FMT may give rise to novel downstream or upstream tarxzsgets to improve lung regeneration and to prevent pulmonary fibrosis.

RNA binding proteins (RBPs) are a kind of proteins that bond to RNA by special motifs (RNA binding domains or RBD), thereby regulating the post-transcriptional modification, transport, decay and translation of RNA ^14^. The role of RBPs is increasingly recognized in development and human diseases including genetic ^15^, neurological ^16^, hepatic ^17^, cardiovascular ^18^, kidney ^19^, malignancy^20^, and, respiratory diseases ^21–24^. Specifically, many RBPs regulate the metabolism of RNA molecules that control the key fibrotic processes such as fibroblast activation, epithelial-mesenchymal transition (EMT) and FMT, thus are implicated in organ fibrosis ^25^.

Human antigen R (HuR) is one of the most widely studied RBPs that possesses multifaceted biological function and the potential as a drug target ^26–28^. HuR protein is encoded by ELAVL1 mRNA. Within the tertiary structure of HuR, there are three RNA recognition motifs (RRMs) that recognize and combine with RNA molecules, and a hinge district that links the RRMs. In pathological circumstances, the perquisite of HuR activation requires a nuclear-cytoplasm shuttle and cytoplasm accumulation mechanism, which is largely regulated through phosphorylation by kinases such as protein kinase C (PKC) and mitogen-activated protein kinase (MAPK) ^29,30^. Such modifications and activation of HuR protein has been found to be a key regulator to organ fibrosis ^25,26,31^. Exogenous TGF-β1 allocates cytoplasmic HuR accumulation that bind to and stabilize TGF-β1 mRNA, which forms a TGF-β1/HuR positive feedback circuit regulating the fibrogenic response in cardiac fibroblasts ^32^. In a mice model of bile duct ligation-induced liver injury and fibrosis, TGF-β1 induced p38MAPK activates HuR through accumulation in the cytoplasm of hepatic cells; HuR silencing reduces hepatic stellate cell activation, oxidative stress, inflammation, collagen and α-SMA expression ^33^. According to another study, HuR is overexpressed in human heart failure; inducible cardiomyocyte-specific HuR-deletion is readily to reduce TGF-β expression, left ventricular hypertrophy, dilation, and fibrosis in a mice model of cardiac hypertrophy ^34^. HuR contributes to Nox4 expression by binding to its 3’-untranslated region (UTR) in hyperglycemia-induced mesangial cell injury and fibrotic kidney remodeling ^35^. Moreover, a recently published study highlights the possible role of HuR in pulmonary fibrosis ^36^. HuR is significantly accumulated in stimulated human lung fibroblasts, IPF lungs and mouse lungs of experimental pulmonary fibrosis; HuR knockdown affected the mRNA stability of ACTA2 ^36^. However, despite numerous studies are favoring the profibrotic role of HuR, recent studies also show that HuR might serve as protective factors against cardiac remodeling and liver fibrosis ^37,38^. Therefore, it still warrants for further investigation about the sophisticated molecular function, the post-translational modification as well as the druggability of HuR in pulmonary fibrosis.

The current study is based on the hypotheses that: 1) HuR is involved in pulmonary fibrosis and TGF-β1-induced FMT via interaction with several key profibrotic RNA molecules, 2) modifications of HuR protein contribute to the functional status of HuR in TGF-β1-induced lung FMT, 3) HuR antagonist CMLD-2 might serve as a promising small molecular agent against FMT and experimental lung fibrosis. We would investigate the expression profiles of key RBPs in FMT by transcriptomics and proteomics, and verify them in fibrotic lung tissues. Then, the impact of HuR overexpression or knockdown on fibroblast activation is studied. HuR binding motifs and HuR-regulated RNA are identified in fibroblasts and TGF-β-transformed myofibroblasts by crosslinking-immunoprecipitation and high-throughput sequencing (CLIP-seq). The effects of CMLD-2 as a HuR antagonist will be further investigated on FMT and on a mice model of bleomycin-induced pulmonary fibrosis.

## 2. Results

### 2.1. Elevated expression of HuR is involved in FMT and lung fibrosis

RNA-seq was performed to identify key regulating genes involving TGF-β1-stimulated fibroblasts *in vitro* (Figure 1). QRT-PCR and Western blot analyses were performed to compare mRNA and protein levels of HuR in IPF lung tissues, rat lung tissues of bleomycin (BLM)-induced pulmonary fibrosis as well as myofibroblastic transdifferentiated from TGF-β1-stimulated fibroblasts. The expressions of HuR gene and protein changed as predicted by the microarray assay and further qRT-PCR investigations, demonstrating that HuR, hnRNPA1, hnRNPE1, TIA1 and TFRC were significantly up-regulated, while QKI5/6, ESRP1, ESRP2 and TTP were significantly down-regulated, while fibroblast activated and acquired myofibroblast markers (including COL1A1 and α-SMA/ACTA2). Co-expression of HuR and α-SMA proteins was also documented by immunofluorescence in both IPF lungs and bleomycin-treated fibrotic rat lung tissues, which supports the role of HuR in FMT in vivo in pulmonary fibrosis. Furthermore, cytoplasmic HuR protein was greatly accumulated while nuclear expression of HuR protein greatly decreased, which indicates the involvement of a nuclear-cytoplasmic shuttle mechanism (Figure 2).

**Figure 1.**
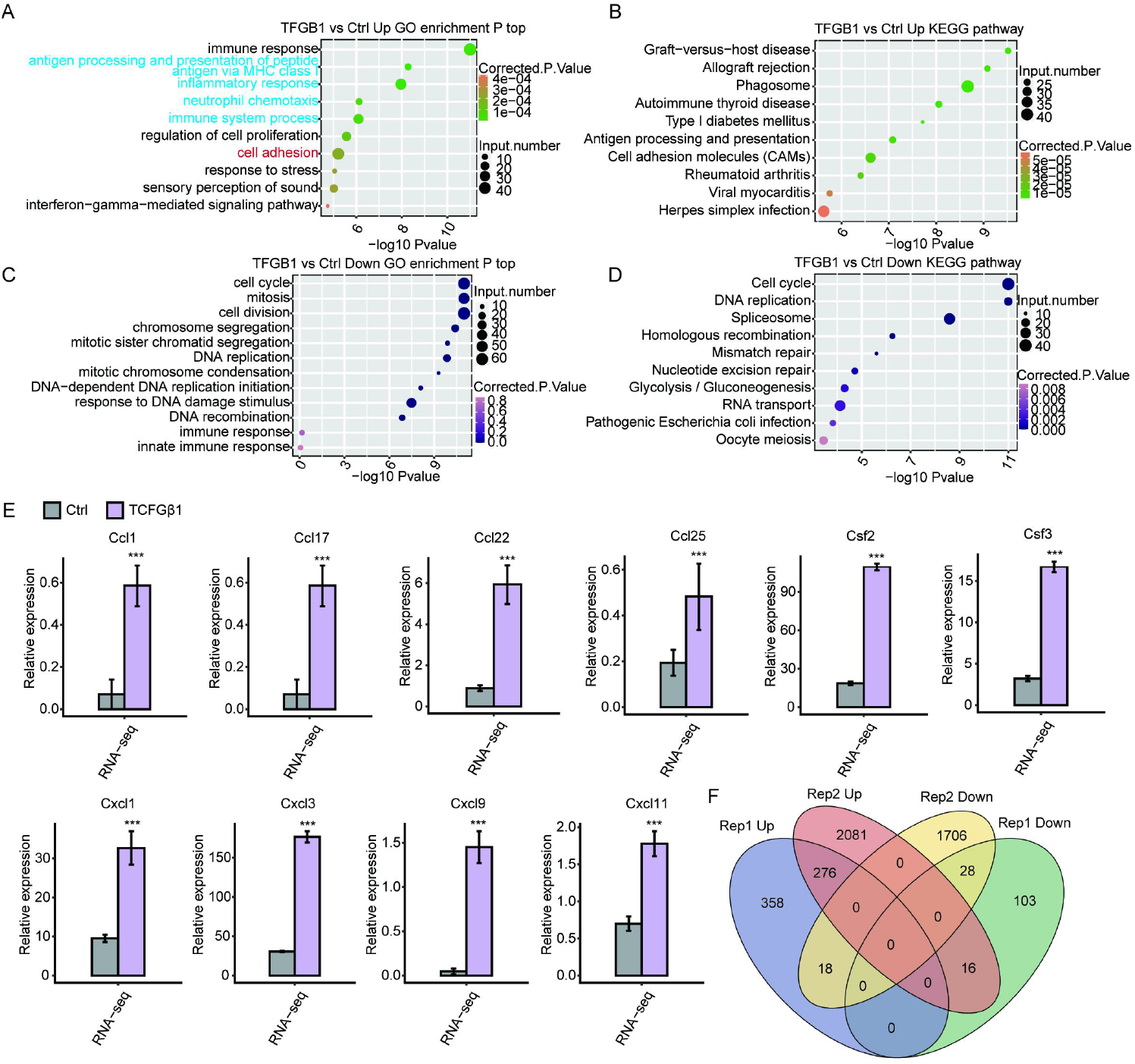
Key regulating genes and cellular events identified by RNA-seq that involves in TGF-β1-stimulated fibroblast-myofibroblast transition *in vitro*.

**Figure 2.**
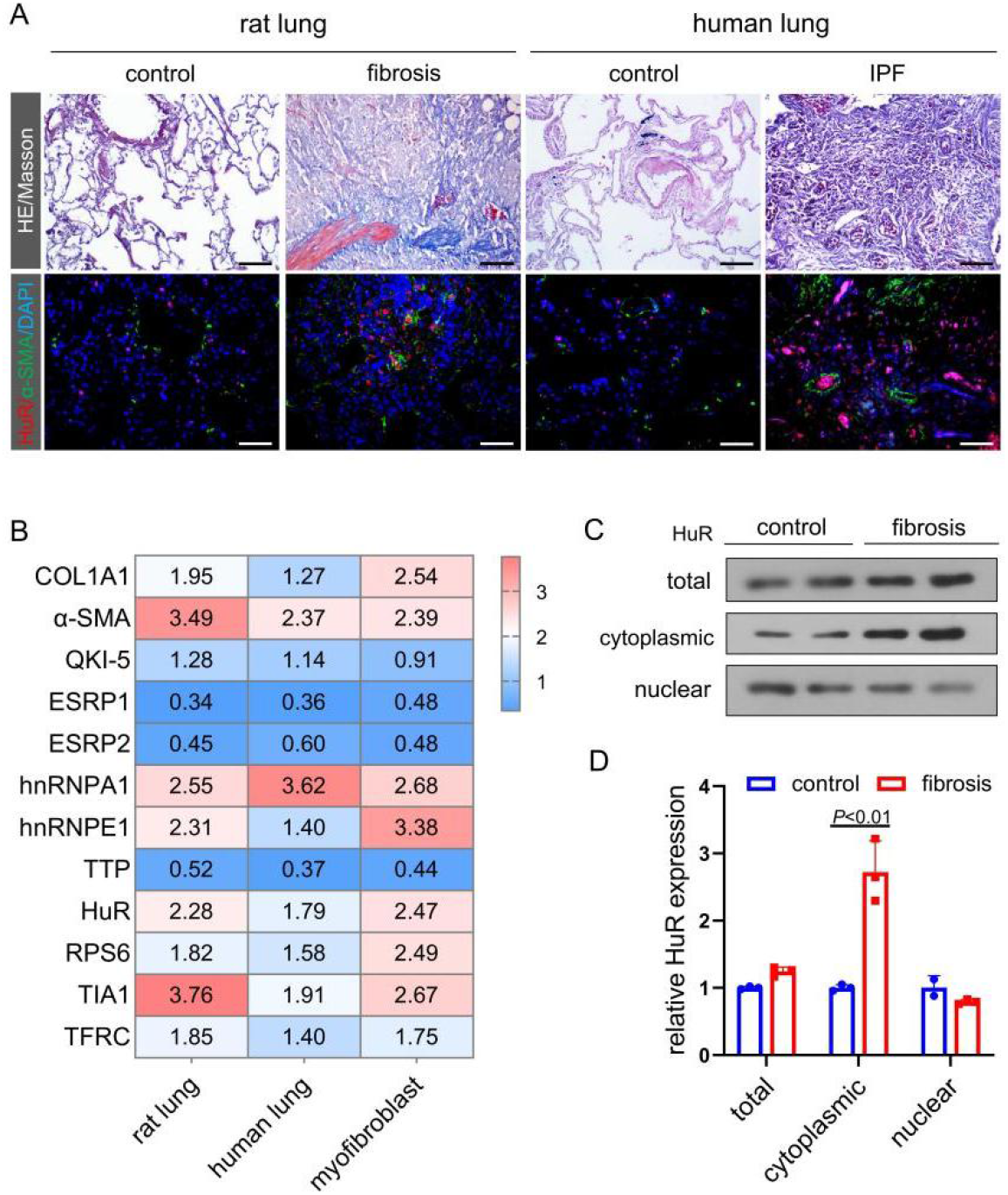
RNA and protein expressions of RBPs, specifically HuR in pulmonary fibrosis and myofibroblast activation. Fig. 1A shows pathology of rat or human lung fibrosis, while enriched cellular immunofluorescence staining of HuR and α-SMA is documented. Fig. 1B shows relative RBP mRNA expressions to GAPDH in rat or lung fibrosis and TGF-β1-stimulated myofibroblasts. HuR mRNA expression is one of the highest among the tested RBPs. Fig 1C & D show the expression of HuR in fibrotic rat lungs, indicating a strong tendency of cytoplasmic accumulation.

### 2.2. HuR overexpression changes fibroblast and myofibroblast biology and genetic expression

To determine the role of HuR in regulating these fate changes in fibroblasts and MFb, we overexpressed HuR in lung fibroblasts and compared the genetic expression coordinates with TGF-β1 syngerically in cell proliferation and cell cycle, while HuR knockdown inhibits TGF-β1-induced fibroblast-myofibroblast transition and activation. Differentially expressed mRNAs were profiled by mRNA-seq in wildtype or HuR-knockdown fibroblasts with or without TGF-β1 stimulation (Figure 3).

**Figure 3.**
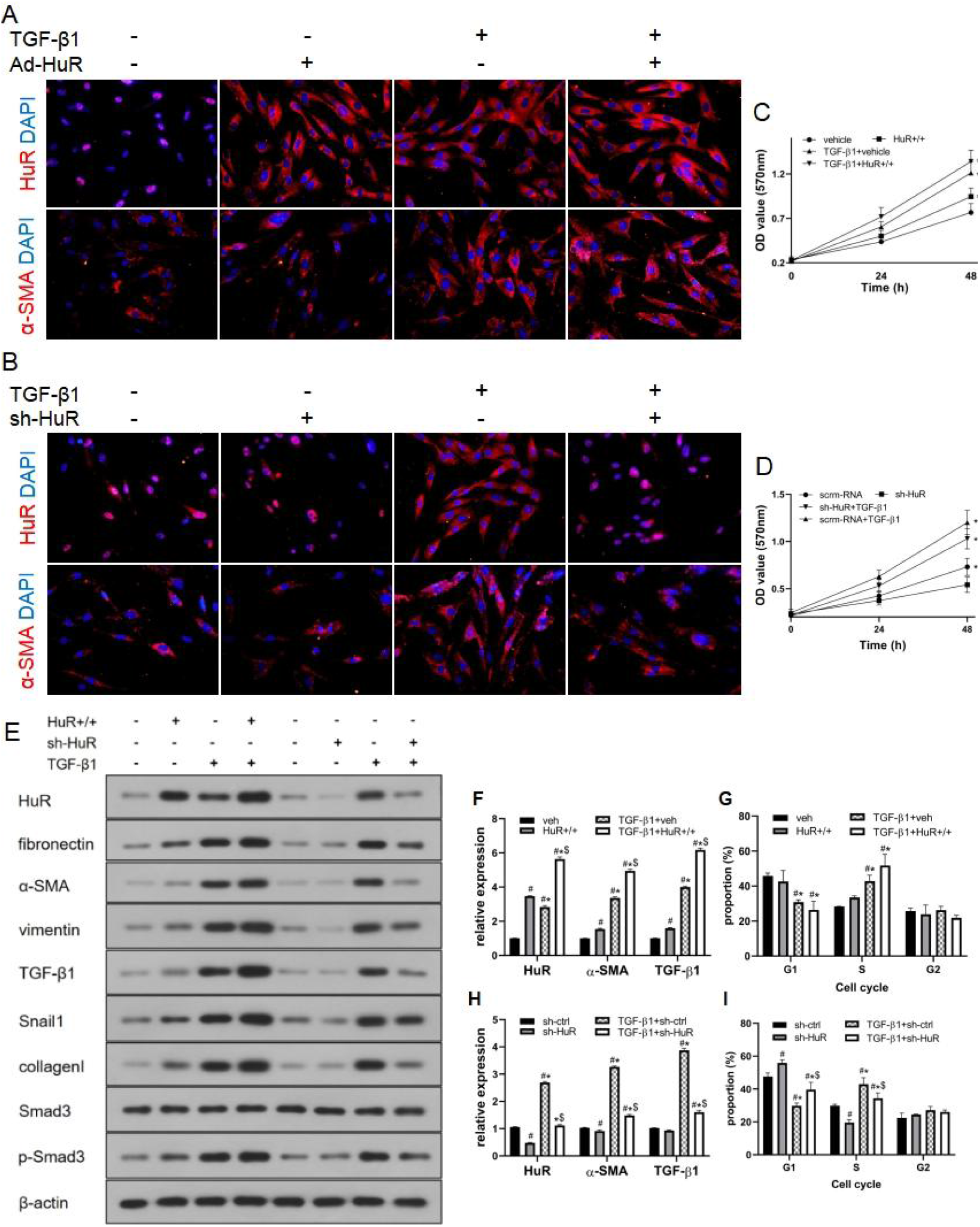
The effect of overexpression (HuR+/+) or knockdown (sh-HuR) of HuR on fibroblast activation and fibroblast-myofibroblast transition. A and B, immunofluorescence. C and D, cell proliferation. E, western blot of key protein expression. G and I, cell cycle analysis.

### 2.3. HuR knockdown inhibits TGF-β1-induced fibroblast activation

We further characterized the role of HuR knockdown in TGF-β1-induced fibroblast-myofibroblast transition. After transfected with Sh-HuR, fibroblasts exhibited less activation status in response to TGF-β1 stimulation. Cell proliferation and myofibroblast marker gene expression was inhibited. Differentially expressed mRNA is profiled in wildtype or HuR-knockdown fibroblasts with or without TGF-β1 stimulation (Figure 3).

### 2.4. HuR-regulated mRNA expression as revealed by CLIP-seq and RNA-seq in TGF-β1-induced fibroblast-myofibroblast transtion

We analyzed the binding motif and RNA of HuR in fibroblasts and myofibroblast (primed by TGF-β1), and performed the GO and KEGG enrichment analysis, as shown here (Figure 4).

**Figure 4.**
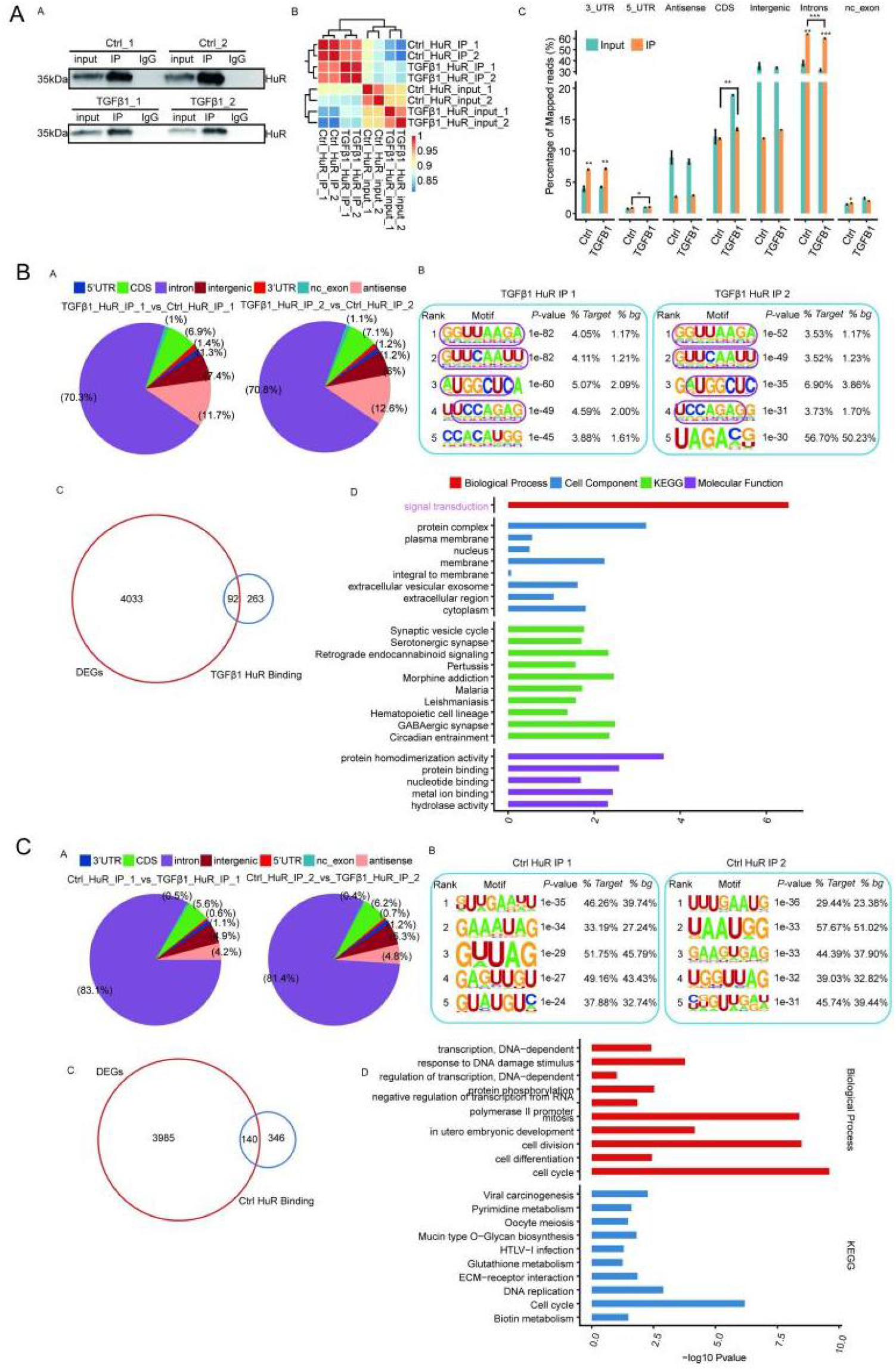
The binding motif and RNA of HuR in fibroblasts and myofibroblast (primed by TGF-β1), and the GO and KEGG enrichment analysis of target genes are shown here.

### 2.5. SPP1 mRNA is stabilized by HuR in TGF-β1-induced fibroblast-myofibroblast transition

We analyzed the candidate key genes that is bond and regulated by HuR. Among them, the HuR specific binding of SPP1, HMBGB2 mRNA was notably elevated and stabilized by stimulation of TGF-β1. After transfection with sh-HuR, fibroblasts exhibited significantly decreased expression of HuR and SPP1 mRNA and proteins. Moreover, we constructed mutant SPP1 plasmids, transfected them into fibroblasts, and evaluated the HuR binding and stability changes of SPP1 mRNA. Since the reporter gene EGFP was connected to the 3’UTR region of SPP1, the fluorescence intensity can well characterize the RNA expression of SPP1. After SPP1 mutant (MT) transfection, SPP1 mRNA increases at 4 and 8 hours, but decreased significantly after 12 and 24 hours. Co-transfection of HuR and SPP1 widetype (WT) groups tended to show stable and slightly higher than that in the SPP1-MT transfection groups, indicating that HuR combined with SPP1 can promote the mRNA stability of SPP1 (Figure 5).

**Figure 5.**
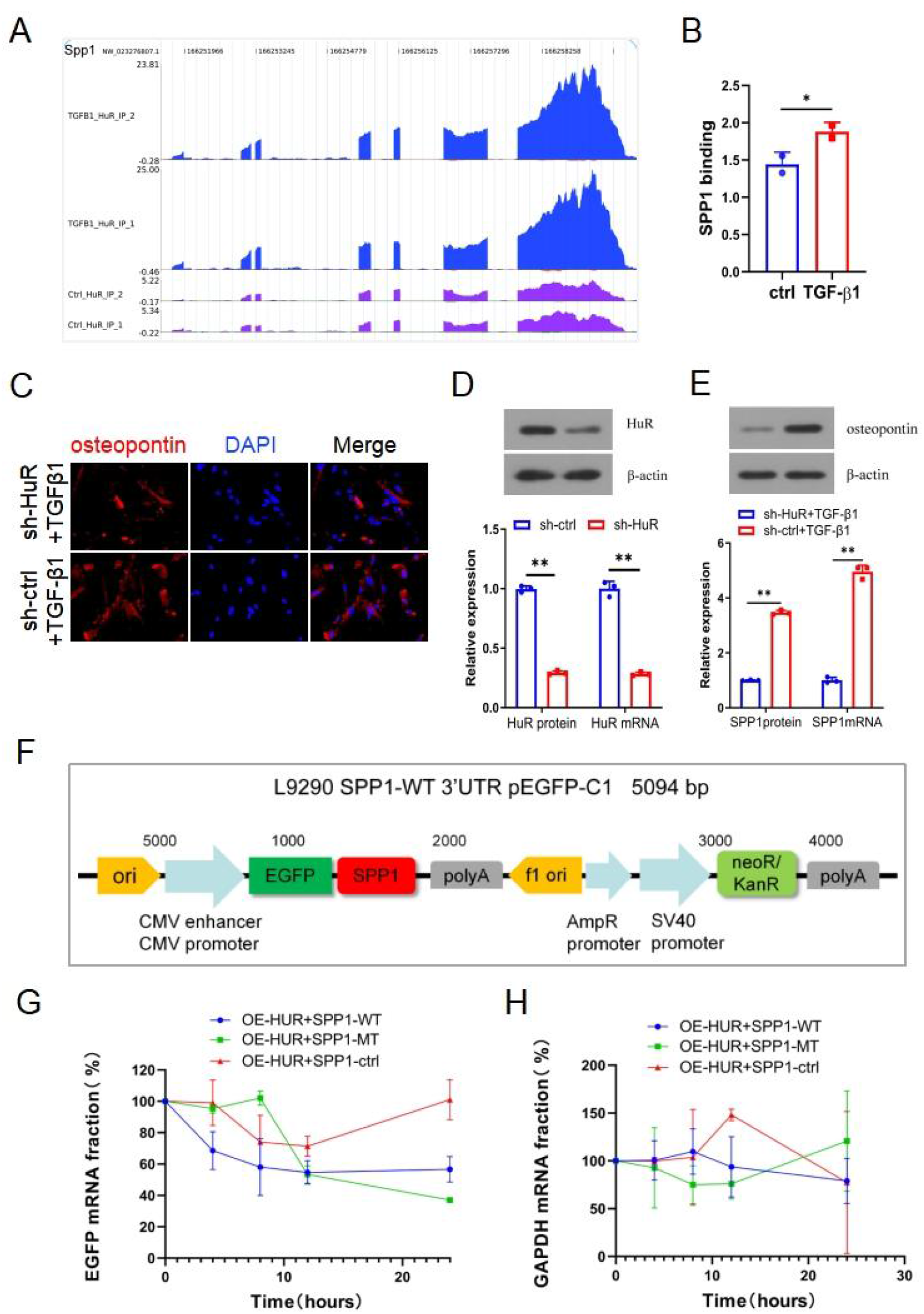
SPP1 is a key target gene of HuR. A and B, binding peaks of SPP1 with HuR. C and D, sh-HuR induces HuR knockdown and inhibits activation of SPP1 gene expression by TGF-β1 in fibroblasts. E, a schematic of the plasmid used for HuR-SPP1 binding. F and G, relative mRNA expression after HuR and SPP1 co-transfection. SPP1-WT (widetype), MT (mutant) and ctrl (control sequence). Co-transfection of HuR and SPP1-ctrl results in stabilized expression of SPP1 mRNA after 8-12 hours.

### 2.6. CMLD-2 inhibits activation and TGF-β1-induced myofibroblast transition of fibroblasts in vitro

The proliferation and motility of fibroblasts were significantly inhibited in the presence of CMLD-2 in a dose-dependent manner. The expression of fibronectin, α-SMA, collagen1, vimentin, Snail1, p-Smad3 was attenuated by CMLD-2. As shown in immunoflurescence staining and Western blot, the expression of cytoplasmic HuR and the ratio of cytoplasmic/nuclear HuR was inhibited by CMLD-2. The inhibitory effect of CMLD-2 on TGF-β1-induced FMT was also seen (Figure 6).

**Figure 6.**
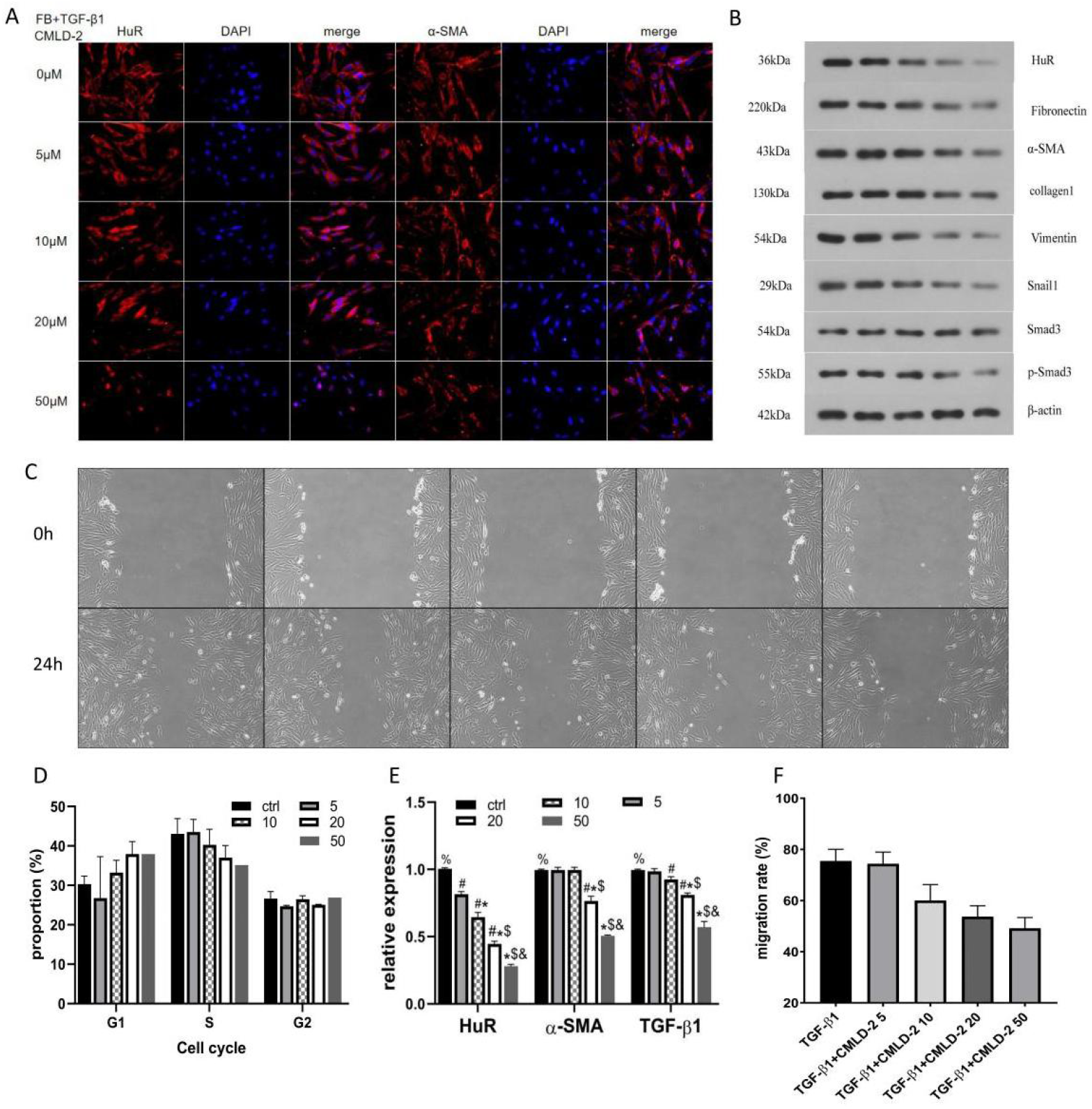
CMLD-2 as an HuR inhibitor shows attenuation of HuR and fibroblast activation markers expression, as well as inhibiting cell migration.

### 2.7. CMLD-2 attenuates bleomycin-induced lung injury and fibrosis in mice

We investigated the impact of CMLD-2 on a mice model of bleomycin-induced lung injury and fibrosis. Introperitoneal injection of CMLD-2 dose-dependently attenuates bleomycin-induced lung injury (Week 2) and fibrosis (Week 4) in mice. The efficacy of CMLD-2 is more prominent of a dose of 50mg/kg/d than 20, 10 and 2 mg/kg/d in attenuating bleomycin-induced lung fibrosis. Consistent is the dose-dependent impact of CMLD-2 on expressions of profibrotic and proinflammatory biomarkers, BALF cytokines and SPP1 expression (Figure 7).

**Figure 7.**
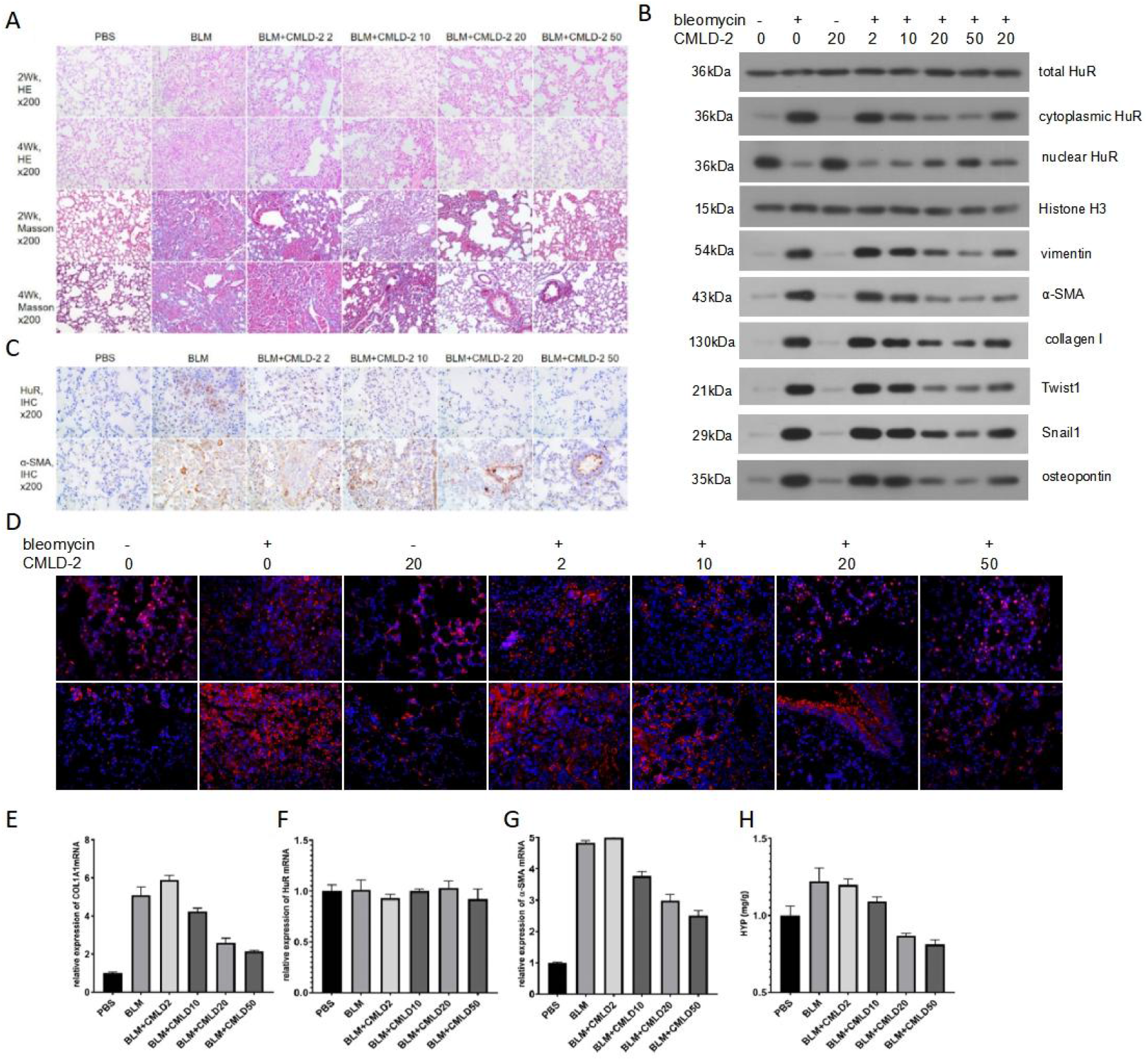
CMLD-2 attenuates bleomycin-induced pulmonary fibrosis in mice. This effect is associated with reduced cytoplasmic HuR accumulation and osteopontin expression, as shown by western blot and immunofluorescence. Lung tissue hydroxyproline levels are also reduced dose-dependently by CMLD-2 in bleomycin-treated mice. D, HuR (upper lane) or osteopontin immunofluorescence in each group.

### 2.8 Upstream activation mechanism of HuR explored by Mass spectrometry and kinase inhibitors

The activity of HuR is suggested to be largely regulated though posttranscriptional modification and kinase-dependent transnuclear shuttle mechanisms. So we first analyzed the phosphorylation loci of HuR in TGF-β1-activated lung fibroblasts. Discordant with our hypothesis however, repetitive MS analysis of HuR co-immunoprecipitation lysate did not reveal any positive phosphorylation site within any of the short peptides (data not shown).

Next we analyzed the effect of PKC or p38MAPK inhibitors (Go-6976 or SB203580, respectively) on subcellular localization and activation of HuR. While TGF-β1 induces transfer of HuR from the nucleus to the cytoplasm, either PKC or p38MAPK inhibition reverses this trend, and attenuate osteopontin mRNA or protein expression. Administration of CMLD-2 also inhibits TGF-β1-induced SPP1 expression, but cytoplasmic HuR accumulation to a less extent (Figure 8).

**Figure 8.**
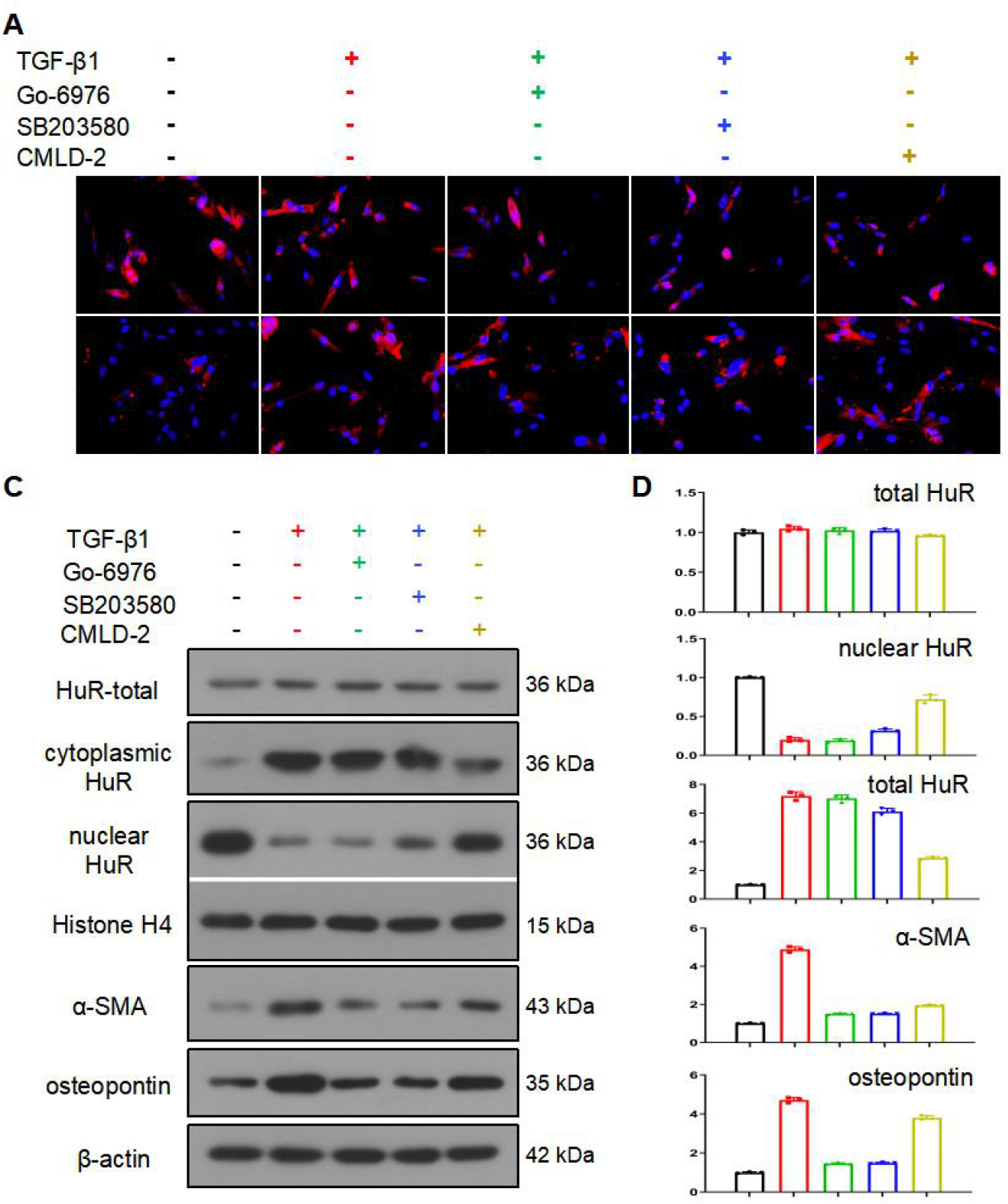
The effects of kinase inhibitors or CMLD-2 on HuR and fibroblast activation. As shown here, CMLD-2, but not Go-6976 or SB203580, significantly alters TGF-β1-induced cytoplasmic accumulation of HuR. However, Go-6976 and SB203580 significantly inhibits osteopontin and α-SMA expression.

## 3. Discussion

Our research tested our hypotheses, and the major findings are as follows: 1) HuR may contribute to pulmonary fibrosis and TGF-β1-induced FMT through modulating SPP1 mRNA metabolism, 2) CMLD-2 exert both *in vivo* and *in vitro* activity of antagonizing HuR and FMT. Our data show that HuR is a profibrotic that is regulated by but not dependent on active TGF-β1 signaling, because HuR knockdown abrogate the effects of TGF-β1 on multiple processes that enable lung fibroblasts to transit into myofibroblasts, while HuR transfection enhances FMT independently or synergically with the presence of TGF-β1. Consequently, migratory capacity, proliferative activity, fibrogenesis, and growth were all significantly inhibited in HuR-depleted fibroblasts as well as myofibroblasts. We also found that lung expression of HuR significantly increases in parallel with myofibroblast accumulation and fibrosis severity in both mouse and rat models of lung fibrosis, in patients with IPF. In addition, HuR is also up-regulated in response to TGF-β1 in cultured lung fibroblast. Among various potential RNA targets, we focus on SPP1 mRNA that codes osteopontin protein. Osteopontin has been identified as a key regulator for activation of fibroblasts and macrophages in pulmonary fibrosis ^39,40^, or as a cellular marker for identify myofibroblast in single cell studies ^41^. The HuR-SPP1 axis is further supported by the effects of CMLD-2 on SPP1 and FMT *in vivo* and *in vitro*. However, this does not exclude other HuR-targeting genes for the universal targeting properties of RBPs. Our study compared the HuR-binding RNA profiles between naïve fibroblasts and TGF-β1-treated fibroblasts; CLIP-seq and RNA half-life data presented here various differently regulated genes covering inflammatory and various cellular processes; the possibly anti-inflammatory effect of CMLD-2 is suggested by its protection from bleomycin-induced early pulmonary injury at day 14. Other previous studies have revealed other downstream genes regulated by HuR in cancers or in fibrotic models, such as proinflammatory genes ^42,43^, Snail1 ^33^ and TGF-β1 ^32,44^. As pulmonary fibrosis is a sophisticated process involving numbers of molecular cell biological mechanisms ^6^, we propose that the universal profibrotic and proinflammatory properties of HuR may justify it as a promising target for fibrosis.

The tertiary structure of HuR protein is composed of three RNA recognition motifs (RRMs) and a hinge motif ^26^. Nuclear-cytoplasm shuttle is primarily dependent on phosphorylation of certain amino acid residues of the hinge motif ^29^. The nuclear-cytoplasm shuttle through posttranslational modification mechanisms, are required in HuR activation in TGF-β1-induced FMT, according to previous studies addressing cellular activation processes in organs other than the lung ^29,33,34, 44–47^. For instance, in rat ventricular myocytes, nuclear-cytoplasmic shuttle of HuR is regulated by SB203580 (a p38MAPK inhibitor) but not by chelerythrine (a PKC inhibitor) ^46^. In liver fibrosis, SB203580 attenuates HuR translocation and activation in bone marrow-derived mesenchymal stem cells ^45^. In ganglions, SB203580 attenuates HuR-stabilizing C/EBP mRNA expression ^47^. In line with these data, we showed here that SB203580 and Go-6976 or CMLD-2 attenuates activation of HuR and SPP1 expression in TGF-β1-primed fibroblasts. However, while SB203580 or Go-6976 inhibits TGF-β1-induced osteopontin and α-SMA expression, these agents exert less effects on cytoplasmic accumulation of HuR, which indicates an HuR-independent mechanism. Moreover, we did not observe any phosphorylation sites in FMT through immunococipitation and MS in all three separate triplicate experiments. The posttranscriptional modulation of HuR through p38MAPK is thus supposed to be organ-nonspecific. While there lacks phosphorylation sites of HuR through Mass spectrometry analysis, phosphorylation of HuR is not dependent on p38MAPK or PKC, or is essential for its activation either. In another aspect, it should be noted that methylation, ubiquitination, transnucleus protein, or CDKs-mediated phosphorylation are otherwise involved in the translocation mechanism of HuR in various cellular models ^29,48^. Such mechanisms are not convincingly excluded as we did not perform here a more comprehensive investigation on these posttranslational mechanisms, although PKC or MAPK is least likely to play major part here. In addition, if we did analyze the phosphorylation status *in vivo* in fibrotic lung tissue, data might be different.

Small molecule antagonists have emerged as an alternative approach in inhibition of HuR functioning ^26,31^. We investigated the efficacy of a small-molecule HuR inhibitor, CMLD-2 ^27,49,50^, against TGF-β1-induced lung fibroblast activation and transformation. Basing on initial pretests, we have determined the effective drug concentration both *in vitro* and *in vivo*, and identified that 20-50 µM CMLD-2 produced obvious inhibitory effect on lung fibroblasts. Using this concentration in subsequent studies, CMLD-2 exhibited preferential inhibitory effects towards fibroblasts and myofibroblasts. CMLD-2-mediated inhibition was accompanied by a G1 phase cell cycle arrest. Molecular studies demonstrated that CMLD-2 treatment significantly reduced HuR mRNA and protein expression, and HuR-regulated profibrotic proteins in fibroblasts and myofibroblasts. Furthermore, our study demonstrate that CMLD-2 is effective against TGF-β1-induced FMT *in vitro* and bleomycin-induced pulmonary fibrosis *in vivo*. While previous studies mainly focus on *in vitro* activity of CMLD-2 ^27,50^, our study revealed the *in vivo* antifibrotic potential of this agent in pulmonary fibrosis, which may open the gate of RBP-targeting small molecules towards lung fibrosis. It’s shown that CMLD-2 slightly reduces the cytoplasmic accumulation of HuR, but more prominently SPP1 expression. This may elicit another question, ie whether CMLD-2 selectively targets HuR and its downstream RNA-modulating mechanisms. As has been reviewed, there emerge a variety of small molecules that targeting HuR fundamentally through various mechanisms ^26,31^. For instance, leptomycin B suppresses the translocation, MS-444 inhibits the dimerization, and CMLD-2 or dihydrotanshinone-I interferes the mRNA binding activities of HuR ^26^. The development of CMLD-2 as a HuR-RNA binding inhibitor was originally established on high-throughput drug screening and reporter RNA techniques ^27^. Additionally, there’s great overlap when comparing our CLIP-seq data with that released by other authors (data not shown) ^51^. As the expression of HuR is greatly activated in FMT or in pulmonary fibrosis, CMLD-2 may exhibit higher efficiency in antagonizing and inhibiting HuR here. Thus, it’s supposed that the effects of CMLD-2 are based basically on its regulation of HuR-RNA binding.

Previous studies have revealed the role of several RBPs in idiopathic pulmonary fibrosis, including cold-inducible RNA-binding protein (CIRBP) that is associated with the prognosis of IPF ^22^, RPS family regulates cellular signaling and activation in fibroblast foci ^24^, or AUF1-Dicer1-miR29-ECM axis ^23^. Excellent studies have elaborated RBP functioning in EMT in cancer cell lines ^25,52–55^. An earlier study by Al-Habeeb^36^ also highlights HuR in pulmonary fibrosis and induction of FMT. These data suggest that RBPs may exert regulating role in FMT. In line with these data, we discovered here that several RBPs including HuR, hnRNPA1, hnRNPE1, TIA1, TFRC, QKI5/6, ESRP1/2 and TTP are differentially expressed in FMT and pulmonary fibrosis as shown by RNA-seq and qRT-PCR. RBP and RNA are targetable, which represents a promising way to inhibit TGF-β1 signaling and fibrosis ^56^. In this scenario, studies are needed to further investigate the role and therapeutic potential of such supposedly fibrosis-regulating RBPs.

Several -omics, RNA-based or RBP-based methods have been utilized to elucidate the role of RBP in regulating RNA processing with regarding to certain cells or diseases. These methods could also be integrated together, including transcriptomics, proteomics, translatomics, RNA interactome capture (RIC), SLAM-seq (to determine the metabolic half-life of bulk RNA), CLIP-seq or RNA pull-down and mass spectrometry (RPD-MS) ^57^. Moreover, except for RNA binding, many RBPs are increasingly be found to take the role of DNA-binding or act as a transcription factor to regulate DNA modification and transcription, which makes their functioning more complicated ^14,15^. Techniques analyzing DNA-protein binding, such as CHIP-seq and ATAC-seq, may be required for analyzing RNA- and DNA-binding RBPs ^15,58^. Despite this, such results should be carefully interpretated, as the regulatory effect of HuR in our study is dependent not on the transcriptional or translational level of HuR expression, but on its posttranslational modification and subcellular redistribution that are regulated by kinases. As shown by RNA-seq and proteomics and further verified by RT-PCR and Western blot, the elevated expression of total HuR protein is not in parallel with that of HuR mRNA in fibrotic lungs or TGF-β1-induced FMT. Since most RBPs are constitutively expressed through various cell types, subcellular proteomic, phosphoproteomics or posttranscriptional modification investigation methods are more precise to determine the modification and functional distribution of RBPs than traditional methods.

There are several shortcomings in our study. Firstly, we did not study the RBPome in fibroblasts or myofibroblasts using RBP screening methods such as RNA interactome capture (RIC) technique ^59^. Such methods are used to identify the sorts of RBPs that are involved in specific cell types or in the process of cell differentiation, either through Oligo d(T) bead capture, click chemistry or phase separation ^60^. In the future we may need to integrate the RIC and multi-omic data to gain a deepened understanding of RBPs that regulates fibrosis and the fate of fibroblasts. Second, RBPs are multi-targeting proteins that may control thousands of RNA molecules. For instance, CLIP-seq data shows that SPP1 is just among one of over 50 thousand HuR-regulating RNA molecules. Additionally, CMLD-2 exert prominent anti-inflammatory effects (eg. Inhibition of cytokine production) otherwise than SPP1 inhibition in the animal model of pulmonary fibrosis. This may obscure the specificity of antifibrotic mechanism of CMLD-2 and druggability of HuR. Third, genetically modified model animals are more specific to elucidate the role and mechanisms of HuR *in vivo* by organ or cell-specific gene overexpression/knockout. Notwithstanding all this, such models have not been adopted in the current study.

Taken together, we disclose a TGF-β1-induced nuclear-cytoplasmic shuttle and activation mechanism of HuR, through which SPP1 mRNA is stabilized to activate fibroblasts in pulmonary fibrosis. The potential of CMLD-2 in attenuating fibroblast-myofibroblast transition and experimental pulmonary fibrosis is also highlighted, possibly through inhibition of TGF-β1-HuR-SPP1 axis. Our study pointed a way through which HuR can be targeted as an RBP by small molecules to inhibit RBP-RNA interactions and key cellular processes in myofibroblast activation and pulmonary fibrosis.

## 4. Materials and Methods

### 4.1. Reagents, materials, specimens and ethical concerns

The current study involved cell culture, animal study and lung specimens obtained from ILD patient donors. All the procedures were in accord with and approved by institutional guidelines for human subject, animal and cell research from the Ethical Committee of Henan Provincial People’s Hospital and Zhengzhou University. The study was conducted according to the guidelines of the Declaration of Helsinki, and approved by the Ethics Committee of Henan Provincial People’s Hospital (protocol code 2018-73, and date of approval December 17th, 2018). Informed consent was obtained from all subjects involved in the study. De-paraffined sections or frozen tissues were collected by bronchoscopic or surgical lung biopsies, or from transplanted lungs with written consents of the patients with a definite multidisciplinary diagnosis of ILD from the Department of Respiratory and Critical Care Medicine, Henan Provincial People’s Hospital. Equipment, reagents and any relevant experimental details are listed in the methods section or detailed in the supplemental materials.

### 4.2. Animal models

Wistar rats or C57/BL6J mice were purchased from and kept in the Experimental Animal Center of Zhengzhou University. Animals got access to water and food *ad libitum*. Rat or mouse model of bleomycin-induced lung fibrosis was induced through intratracheal injection of bleomycin solution (5mg/kg for mice, 15mg/kg for rats; CAS No. 9041-93-4, Aladdin Biochemical Technology Co. Ltd., Shanghai, China). CMLD-2 was dissolved in dimethyl sulfoxide (DMSO) and injected intraperitoneally once daily, 5 days per week. The doses of CMLD-2 (0-50mg/kg/d) were determined by referring to previous studies and preliminary experiments. Animals were sacrificed at different time points under general anaesthesia with intraperitoneal pentobarbital sodium. Lung sample preparation, bronchoalveolar lavage (BAL) and BAL fluid cell count was performed and as previously described ^61^.

### 4.3. Cell culture

MRC-5 lung fibroblast and 293T epithelial cell lines (purchased from National Collection of Authenticated Cell Cultures, Shanghai, China) were cultured in RPMI1640 culture medium supplemented with 10% fetal bovine serum (FBS, Gibco, Life Technologies) and 1% penicillin/streptomycin (Gibco, Life Technologies) at 37°C in a humidified atmosphere containing 5% CO2 for 72h. Cells were cultured to 70∼80% confluency, and starved in 0.2% FBS-containing medium. After 24 hours, cells were incubated with human recombinant TGF-β1 (5 ng/ml, 10804-HNAC, Sino Biological Inc., Beijing, China), CMLD-2 (LN03203514, Wuxi Apptec, Wuxi, China), Go-6976 (10 mM, a PKC inhibitor, G276069, Aladdin Biochemistry Co. Ltd., Shanghai, China) or SB203580 (10 mM, a p38MAPK inhibitor, S131899, Aladdin Biochemistry Co. Ltd., Shanghai, China) with different incubation concentrations for 48h. Cellular protein or total RNA was then harvested and analyzed.

### 4.4. HuR plasmid or shRNA transfection

Cells were raised at a 70% confluence, and transfected with plasmid or shRNA using Lipofectamine 2000 (11668-019, Invitrogen) and opti-MEM (31985070, Invitrogen) following the manufacturer’s instructions. For each 6-well plate, solution 1 was prepared by mixing 100 μL opti-MEM and 6 μL Lipofectamine 2000, and place at room temperature for 5min. Solution 2 was a mixture of opti-MEM 100 μL and 2 μg plasmid. Solutions 1 and 2 were mixed and incubated at room temperature for 20min, then added into the cells slowly and shake gently. The cells were cultured 24 h for RT-PCR or 48 h for Western blot detection, respectively. Transfection efficiency was determined through RT-PCR or Western blot assay. The sequences and maps of HuR plasmid and shRNAs are listed in the supplemental methods.

### 4.5. Cell proliferation assay

Cell proliferation was measured with a MTT method. Briefly, cells were plated at a density of 3×10^3^ cells per well in 96-well plates, with the indicated treatment and time. MTT (20μl) was added per well to the culture medium for a further incubation period of 4 h. Then the OD values were measured under 570nm with ELX-800 (BioTek, VT, USA).

### 4.6. Scratch assay

Cells were kept in serum-free medium to produce a confluent monolayer and treated with mitomycin C (1μg/ml) for 1 h. Scratch was induced by 200 μL pipette tip. Photos were taken at 0h and 24 hours, and then the width of the scratch was analyzed by Image J software (version 1.52v, http://imagej.nih.gov/ij/).

### 4.7. Flow cytometry

Cell cycle was determined through flow cytometry. Cells were re-suspended with a concentration of 1×10^6^/ml, centrifuged at 1000g for 5 min. Cold ethanol (1ml, 70%) was added and then incubated at 4℃ overnight. Cells were centrifuged and washed by pre-cool PBS twice. 100μl RNase A was added, then the material was succumbed to water bath (37℃) for 30 min. Staining was performed at 4℃ in dark room after 500μl propidium iodide was added. Specimens were then transferred to a NovoCyte Flow Cytometry Analyzer (Aceabio, CA, USA) for analysis.

### 4.8. Histology, immunohistochemistry and immunofluorescence

Lung sections (5 μm thickness) were prepared from paraffin-embedded tissue specimens and then subjected to conventional hematoxylin and eosin staining (HE) and Masson’s trichrome. IHC or IF staining was performed on deparaffinized lung sections and/or cell samples as described before ^61^. HE/Masson/IHC or IF images were acquired using a BX53 fluorescence microscope (Olympus, Japan).

### 4.9. Enzyme linked immunosorbent assay (ELISA)

Indirect ELISA was performed on BALF samples to quantify cytokine production using commercially available kits following the instructions. TGF-β1 (SEA124Mu), IL-4 (SEA077Mu) and IL-10 (SEA056Mu) ELISA kits were purchased from Cloud Clone Corp., Wuhan, China. TNF-α (EMC102a.96) and IL-1β (EMC001b.48) ELISA kits were purchased from Neobioscience Biotechnique Co. Ltd., Shenzhen, China.

### 4.10. Hydroxyproline assay

Hydroxyproline assay was performed of lung tissue samples using a hydroxyproline assay kit (Nanjing Jiancheng Bioengineering Institute, Nanjing, China) according to the instructions.

### 4.11. Western blot

Western blot was performed as described before ^61^. Total protein was extracted from cells or -80°C stored lung tissue samples and quantified using a bicinchonininc acid (BSA) protein quantification kit. Proteins were then separated by SDS-PAGE (sodium dodecyl sulfate polyacrylamide gel electrophoresis) and transferred to PVDF (polyvinylidene difluoride) membranes. After block, primary and second antibody staining, membranes were subjected to chemiluminescence (SuperSignal West, Thermo). Band intensities were analyzed using Gel-Pro-Analyzer software (Media Cybernetics, MD, USA).

### 4.12. Quantitative real-time PCR (qRT-PCR)

Total RNA was extracted from cells or lung tissue samples that had been stored at -80°C with TRIZOL reagent ((Thermo, USA) and chloroform. RNA quality and concentration was measured by a NANO 2000 spectrophotometer. Reverse transcription was performed with a PCR instrument (Exicycler 96, BIONEER, Korea) as follows: adding to RNA specimens with oligo (dT)15 1 μl and random 1 μl, and supplement with ddH2O to allow a total volume of 12.5 μl; then incubate at 70℃ for 5 min and ice-bath for 2 min. dNTP (2.5 mM each) 2 μl, 5×Buffer 4 μl, Rnase inhibitor 0.5 μl and M-MLV (1 μl or 200 U) was added to the supernatant and gently mixed after a short period of centrifuge, then subjected to 25℃ for 10 min, 42℃ for 50 min, and finally 80℃ for 10 min to terminate the reaction. cDNA was harvested for further procedures. The reaction system of real-time PCR was: cDNA template 1 μl, forward primer (10 μM) 0.5 μL, reverse primer (10 μM) 0.5 μL, SYBR GREEN mastermix 10 μl, ddH2O (add to a final total volume of 20 μL). Incubation cycle was: 94℃ for 5 min, 94℃ for 10 s, 60℃ for 20 s, 72℃ for 30 s; scan and go to line 2, cycle 40: incubate at 72℃ for 2 min 30s, incubate at 40℃ for 1 min 30 s, melting 60℃ to 94℃, 1 s per every 1.0℃, incubate at 25℃ for 1-2 min. RNA content was analyzed basing on the Ct method compared to the reference gene.

### 4.13. RNA stability assessment

Cells were treated with 2 mM actinomycin D and CMLD-2 (or DMSO as a control). RNA was extracted at several time points (0, 30, 60, 90min) and quantified by qRT-PCR for mRNA expression.

### 4.14. RNA-seq analysis

Total RNA was treated with RQ1 DNase (Promega) to remove DNA. The quality and quantity of the purified RNA were determined by measuring the absorbance at 260nm/280nm (A260/A280) using smartspec plus (BioRad). RNA integrity was further verified by 1.5% Agarose gel electrophoresis. For each sample, 1μg of total RNA was used for RNA-seq library preparation. Polyadenylated mRNAs were purified and concentrated with oligo(dT)-conjugated magnetic beads (Invitrogen) before being used for directional RNA-seq library preparation. Purified mRNAs were fragmented at 95 ℃ in fragmentation buffer followed by end repair and 5’ adaptor ligation. Then reverse transcription was performed with RT primer harboring 3’ adaptor sequence and randomized hexamer. The cDNAs were purified and amplified and PCR products corresponding to 200-500 bps were purified, quantified, and stored at -80 ℃ before sequencing. For high-throughput sequencing, the libraries were prepared following the manufacturer’s instructions and applied to Illumina HiSeq X Ten system for 150 nt paired-end sequencing.

Raw reads containing more than 2-N bases were first discarded. Then adaptors and low-quality bases were trimmed from raw sequencing reads using FASTX-Toolkit (Version 0.0.13). The short reads less than 16nt were also dropped. After that, clean reads were aligned to the GRch38 genome by tophat2 allowing 4 mismatches ^62^. Uniquely mapped reads were used for gene reads number counting and FPKM calculation (fragments per kilobase of transcript per million fragments mapped) ^63^. The R Bioconductor package edgeR was utilized to screen out the differentially expressed genes (DEGs) ^64^. A false discovery rate (FDR) <0.05 and fold change >2 or < 0.5 were set as the cut-off criteria for identifying DEGs. All DEGs were annotated to the Gene Ontology database and the Kyoto Encyclopedia of Genes and Genomes (KEGG) pathway database.

The alternative splicing events (ASEs) and regulated alternative splicing events (RASEs) between the samples were defined and quantified as described previously ^65^. To elucidate the validity of the RNA-seq and ASE data, qRT-PCR was performed for selected DEGs.

### 4.15. RNA binding protein immunoprecipitation (RIP), RIP-PCR and CLIP-seq analysis

For HuR protein-RNA co-immunoprecipitation, cells were first lysed in ice-cold lysis buffer (1×PBS, 0.5% sodium deoxycholate, 0.1% SDS, 0.5% NP40) with RNase inhibitor (Takara, 2313) and a protease inhibitor (Solarbio, 329-98-6) on ice for 5 min. The mixture was then vibrated vigorously and centrifuged at 13,000 x g at 4℃ for 20 min to remove cell debris. The supernatant was incubated with DynaBeads protein A/G (Thermo Fisher, 26162) conjugated with anti-HuR antibody (Proteintech, 11910-1-AP, Wuhan, China) or normal IgG at 4℃ overnight. The beads were washed with Low-salt Wash buffer, High-salt Wash buffer and 1× PNK Buffer, respectively. Resuspend the beads in Elution Buffer and then divided into two groups, one for RNA isolation from HuR-RNA complexes and another for the western blotting assay for HuR. The HuR-bound RNAs were isolated from the immunoprecipitation of anti-HuR using TRIzol (Invitrogen). For RIP-PCR analysis, specific RNA primers were used in qRT-PCR to identify and quantify HuR-targeting RNA.

For CLIP-seq library preparation and sequencing, complementary DNA (cDNA) libraries were prepared with the KAPA RNA Hyper Prep Kit (KAPA, KK8541) according the manufacturer’s procedure and high-throughput sequencing of the cDNA libraries was performed on an Illumina Xten platform for 150 bp paired-end sequencing. After reads were aligned onto the genome with TopHat 2 ^62^, only uniquely mapped reads were used for the following analysis. ABLIRC strategy was used to identify the binding regions of HuR on genome ^65^. Reads with at least 1 bp overlap were clustered as peaks. For each gene, computational simulation was used to randomly generated reads with the same number and lengths as reads in peaks. The outputting reads were further mapped to the same genes to generate random max peak height from overlapping reads. The whole process was repeated for 500 times. All the observed peaks with heights higher than those of random max peaks (p-value < 0.05) were selected. The IP and input samples were analyzed by the simulation independently, and the IP peaks that have overlap with Input peaks were removed. The target genes of IP were finally determined by the peaks and the binding motifs of IP protein were called by HOMER software ^66^. To sort out functional categories of peak associated genes (target genes), Gene Ontology (GO) terms and KEGG pathways were identified using KOBAS 2.0 server ^67^. Hypergeometric test and Benjamini-Hochberg FDR controlling procedure were used to define the enrichment of each term.

### 4.16. SPP1 mutant construction, transfection and mRNA stability assay

To confirm the effect of HuR binding on SPP1 mRNA metabolism, we construct the mutant SPP1 construct and cloned them with EGFP gene, and then co-transfected cells with HuR and widetype (WT) or mutant (MT) SPP1 plasmids. Cells were seeded by 2×10^5^ and cultured overnight. JetPRIME buffer (75µl), plasmid (0.8 μg) and JetPRIME reagent (1.6µl) (PT-114-75, Polyplus, Illkirch, FRANCE) were mixed and added to the culture medium to allow incubation for 4 hours. After 48 hours of transfection, the successfully transfected cells were treated with actinomycin D (10 µg/ml), and the cells were collected at five time points (0, 4, 8, 12 and 24 hours) to extract total RNA. The stability of EGFP gene characterized the regulation of HuR gene on the binding stability of SPP1 gene segment. GAPDH is the control detection gene and actin is the internal reference gene. Transfection efficiency and mRNA stability was analyzed by qRT-PCR. cDNA synthesis was done by standard procedures and RT-qPCR was performed on the Bio-Rad S1000 with Hieff qPCR SYBR® Green Master Mix (Low Rox Plus; YEASEN, China). The concentration of each transcript was then normalized to GAPDH mRNA level using 2-ΔΔCT method. The information of primers is presented as follows. Elavl1-F: CATCTTCATCTACAACCTTGG. Elavl1-R: GCCGTTCAGACTTGCTAT. EGFP-F: CCGACCACTACCAGCAGAA. EGFP-R: CGAACTCCAGCAGGACCAT. ACTB-F1: GAGAAAATCTGGCACCACACC. ACTB-R1: GGATAGCACAGCCTGGATAGCAA. GAPDH-F: GGTCGGAGTCAACGGATTTG. GAPDH-R: GGAAGATGGTGATGGGATTTC.

### 4.17. Statistical analysis

Data were expressed as ratio or mean ± standard deviation (SD), and analyzed through Graphpad Prism software version 8.3.0 (GraphPad Software, CA, USA), R software version 4.0.5 (https://www.r-project.org/) or SPSS package version 26.0 (IBM Inc., NY, USA). Statistical differences were determined by Student’s t-test, χ^2^ test or one-way ANOVA, followed by post-hoc SNK test. *P* values less than 0.05 were considered to be statistically significant.

## Supplementary Materials

The following are available online at the current website.

## Author Contributions

G.Q., R.H., L.M., L.L., L.W., Z.J., W.X., H.Y., Z.K., A.Y. and L.L. performed experiments. W.Z., X.Z. and W.Z. performed experiments and analyzed the data. W.L. collected lung biopsy samples. G.Z., U.C., W.Z. and Z.X drafted the manuscript. All authors have read and agreed to the published version of the manuscript.

## Funding

This research was funded by the National Natural Science Foundation of China (grant No. 81600047), the Natural Science Foundation of Henan Province (grants No. 142300410381, 142102310409), Zhongyuan Thousand Talent Program-Youth Outstanding Talent (grant No. 2018), Research Program of the Education Department of Henan Province (grant No. 21A320001), Research Program of the Health Commission of Henan Province (grants No. LHGJ20190581, YXKC2020043), Zhongyuan Thousand Talent Program-Science and Technology Innovation Leader (grant No. 2018), Henan Provincial Major Project of Public Welfare (grant No. 201300310500).

## Acknowledgments

We would thank Dr. Chen Wen (Wuhan ABLife Inc., Wuhan, China) for her technical help and valuable suggestions on drafting this manuscript.

## Conflicts of Interest

The authors declare no conflict of interest. The funders had no role in the design of the study; in the collection, analyses, or interpretation of data; in the writing of the manuscript, or in the decision to publish the results.

## Abbreviations

ARE: AU-rich element
BALF: Bronchoalveolar lavage fluid
BLM: Bleomycin
CLIP-seq: Crosslinking-immunoprecipitation and high-throughput sequencing
ECM: Extracellular matrix
EMT: Epithelial-mesenchymal transition
FMT: Fibroblast-myofibroblast transition
ILD: Interstitial lung disease
IPF: Idiopathic pulmonary fibrosis
MFb: Myofibroblast
MS: Mass spectrometry
P38MAPK: P38 mitogen-activated protein kinase
PKC: Protein kinase C
RBD: RNA binding domain
RBP: RNA binding protein
RIC: RNA interactome capture
RIP: RNA binding protein immunoprecipitation
RPD: RNA pull-down
RRM: RNA recognition motif
TGF-β: Transforming growth factor β
UTR: Untranslated region

---

**Supplemental method 1.**
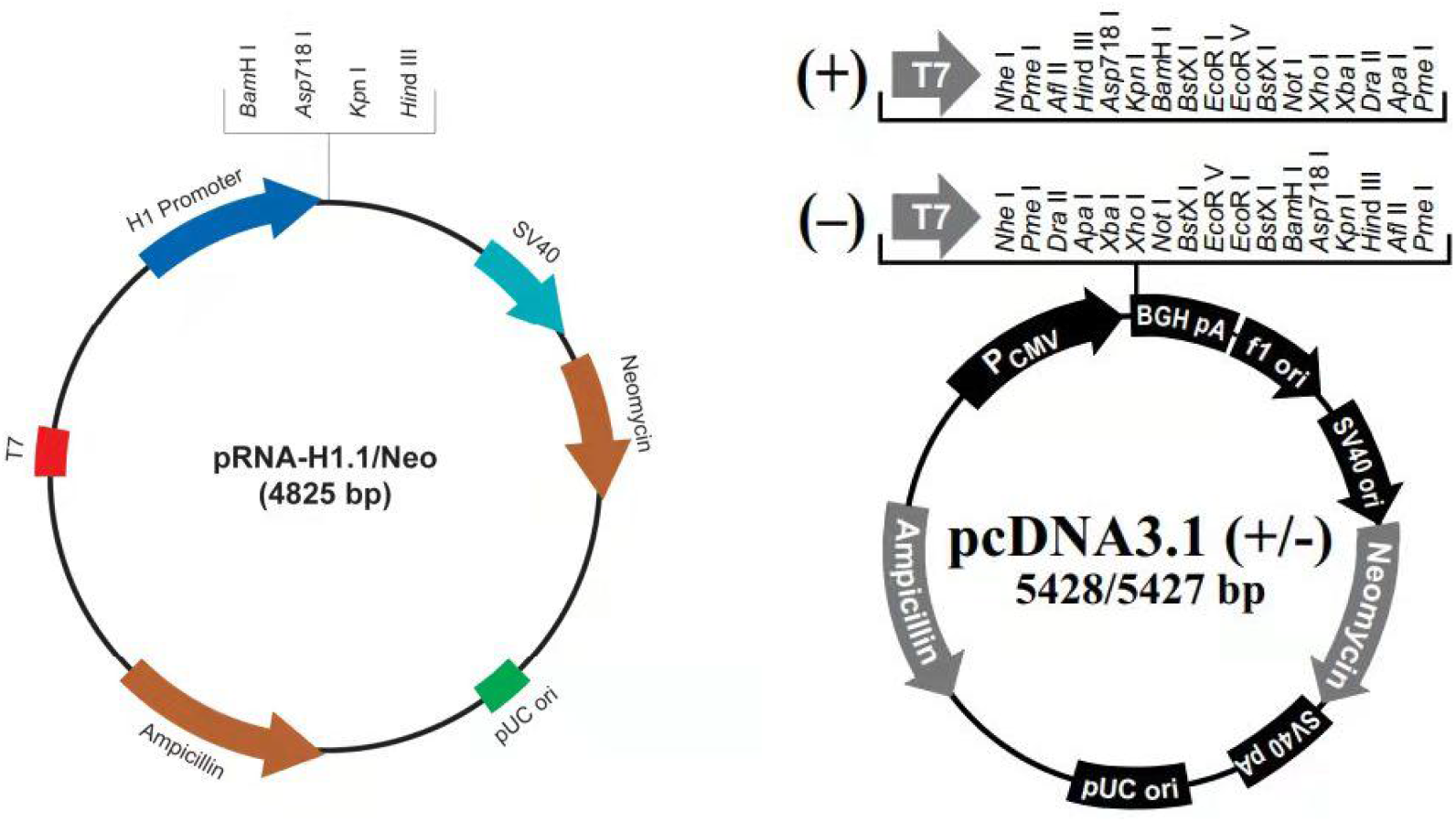
The sequences of sh-HuR and HuR plasmid and their maps. sh-HuR1 : gGCCCATAGCTAGCATTCAACAttcaagagaTGTTGAATGCTAGCTATGGGCttttt; sh-HuR2 : gGGTTTGGCTTTGTGACCATGAttcaagagaTCATGGTCACAAAGCCAAACCttttt; sh-HuR3: gGGTTTGGGCGGATCATCAACTttcaagagaAGTTGATGATCCGCCCAAACCttttt. HuR plasmid CDS (981bp): ATGTCTAATGGTTATGAAGACCACATGGCCGAAGACTGCAGGGGTGACATCGGGAGAACGAATTTGATCGTCAACTACCTCCCTCAGAACATGACCCAGGATGAGTTACGAAGCCTGTTCAGCAGCATTGGTGAAGTTGAATCTGCAAAACTTATTCGGGATAAAGTAGCAGGACACAGCTTGGGCTATGGCTTTGTGAACTACGTGACCGCGAAGGATGCAGAGAGAGCGATCAACACGCTGAACGGCTTGAGGCTCCAGTCAAAAACCATTAAGGTGTCGTATGCTCGCCCGAGCTCAGAGGTGATCAAAGACGCCAACTTGTACATCAGCGGGCTCCCGCGGACCATGACCCAGAAGGACGTAGAAGACATGTTCTCTCGGTTTGGGCGGATCATCAACTCGCGGGTCCTCGTGGATCAGACTACAGGTTTGTCCAGAGGGGTTGCGTTTATCCGGTTTGACAAACGGTCGGAGGCAGAAGAGGCAATTACCAGTTTCAATGGTCATAAACCCCCAGGTTCCTCTGAGCCCATCACAGTGAAGTTTGCAGCCAACCCCAACCAGAACAAAAACGTGGCACTCCTCTCGCAGCTGTACCACTCGCCAGCGCGACGGTTCGGAGGCCCCGTTCACCACCAGGCGCAGAGATTCAGGTTCTCCCCCATGGGCGTCGATCACATGAGCGGGCTCTCTGGCGTCAACGTGCCAGGAAACGCCTCCTCCGGCTGGTGCATTTTCATCTACAACCTGGGGCAGGATGCCGACGAGGGGATCCTCTGGCAGATGTTTGGGCCGTTTGGTGCCGTCACCAATGTGAAAGTGATCCGCGACTTCAACACCAACAAGTGCAAAGGGTTTGGCTTTGTGACCATGACAAACTATGAAGAAGCCGCGATGGCCATAGCCAGCCTGAACGGCTACCGCCTGGGGGACAAAATCTTACAGGTTTCCTTCAAAACCAACAAGTCCCACAAATAA All the shRNAs are of 57bp, linked to pRNAH1.1 vector, and the restriction enzyme cutting sites are BamHI/HindIII. The HuR plasmid sequence is linked to pcDNA3.1 vector. The restriction enzyme sites are HindIII/XhoI.

**Supplemental method 2.**
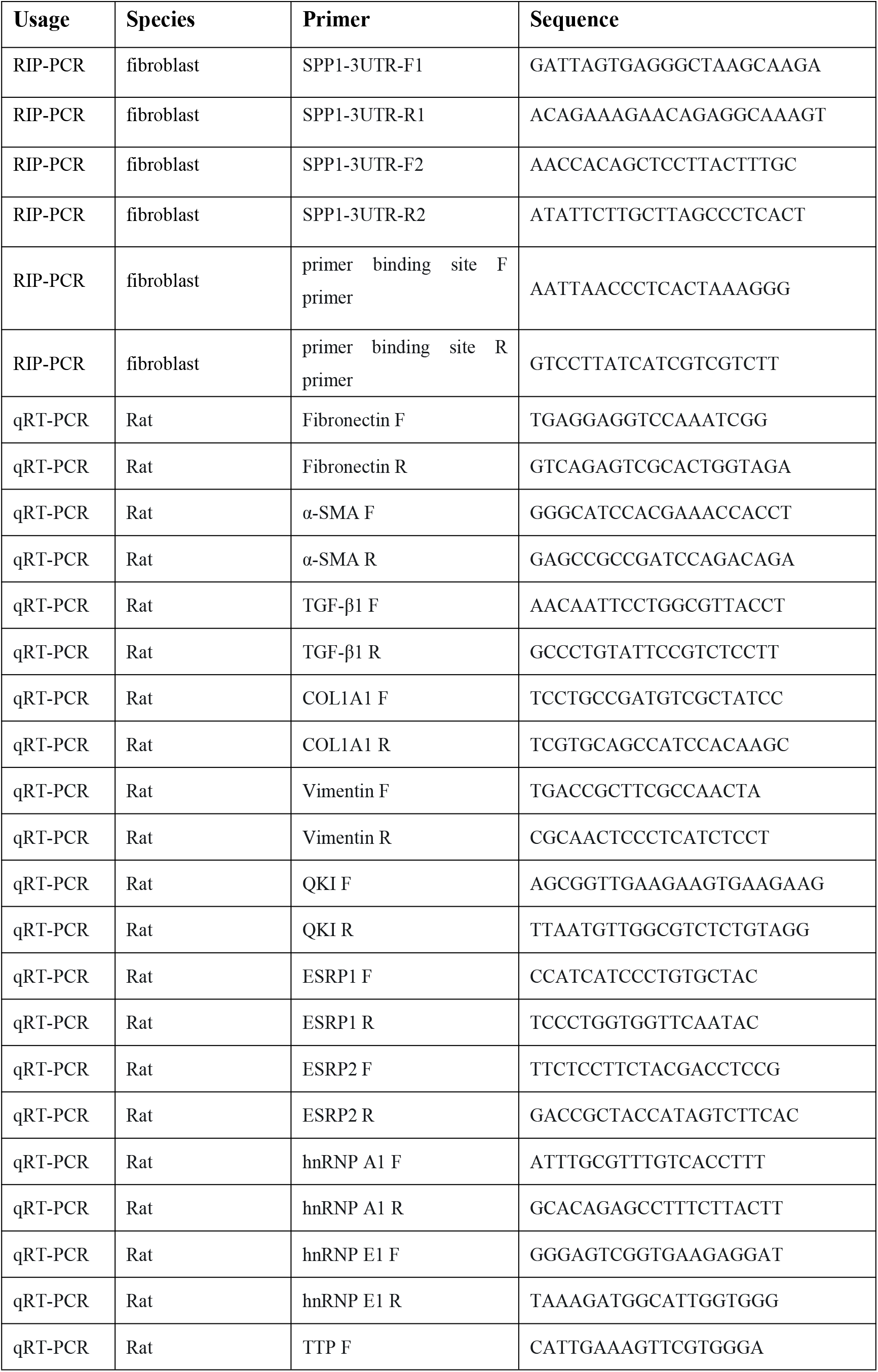
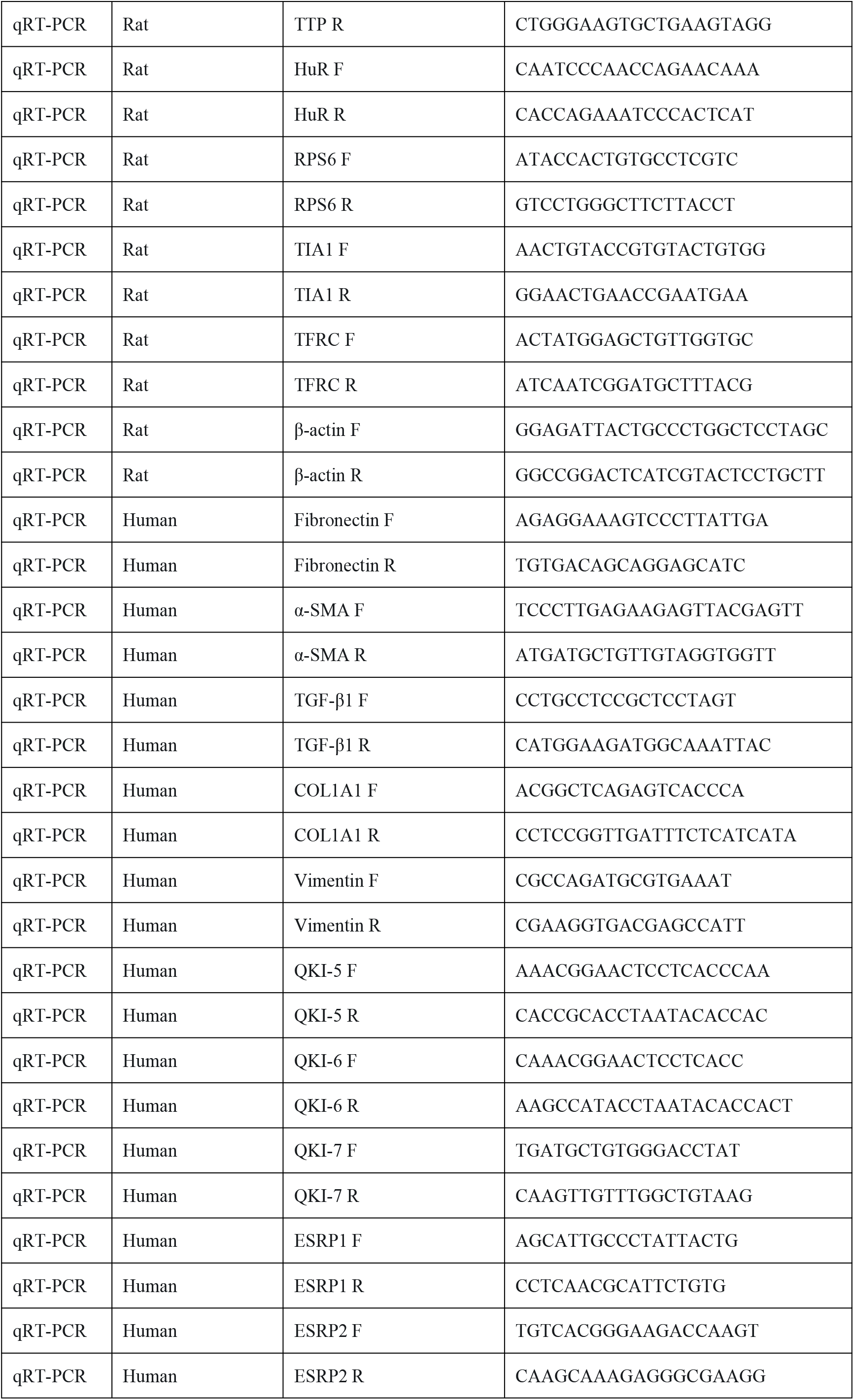
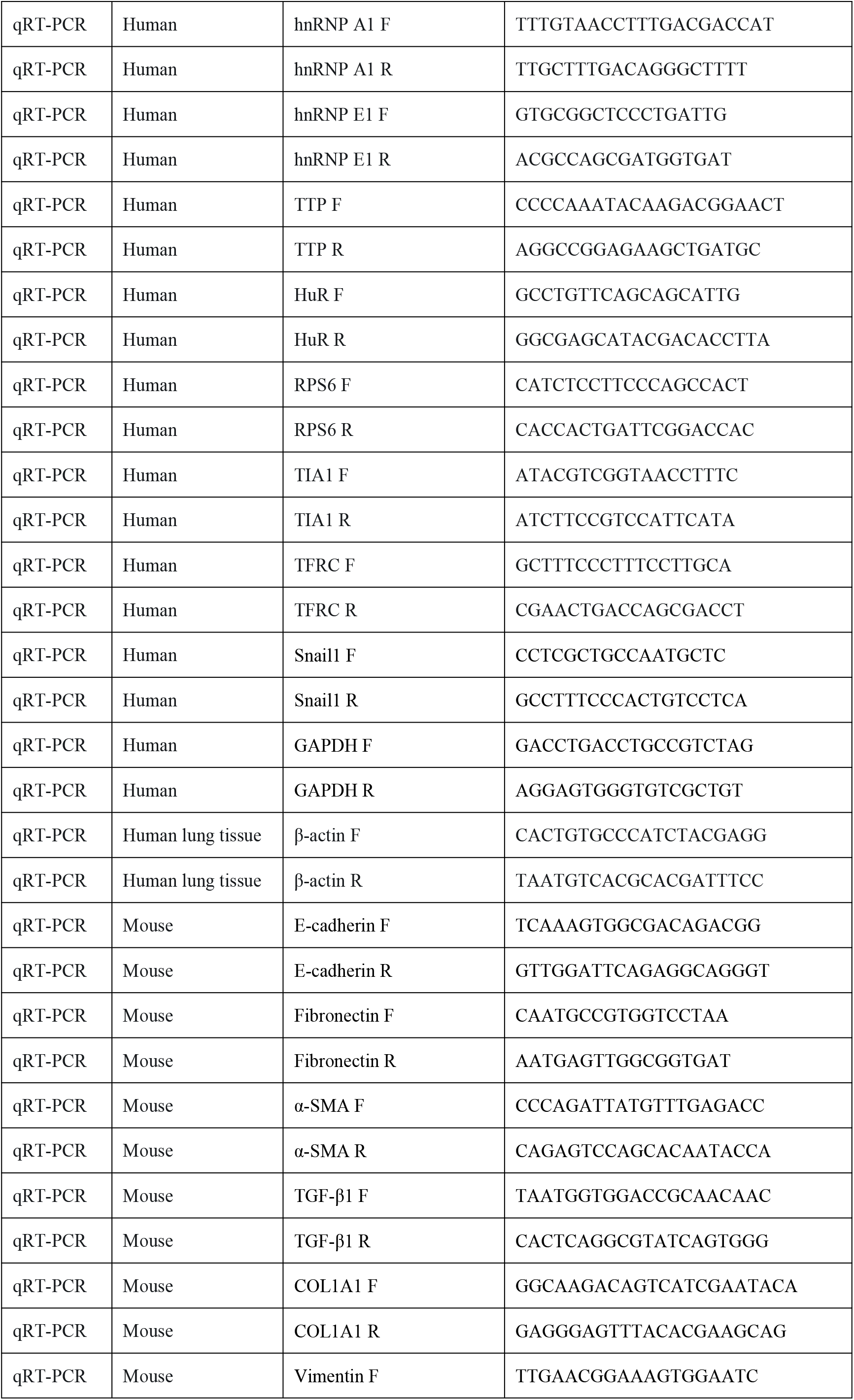
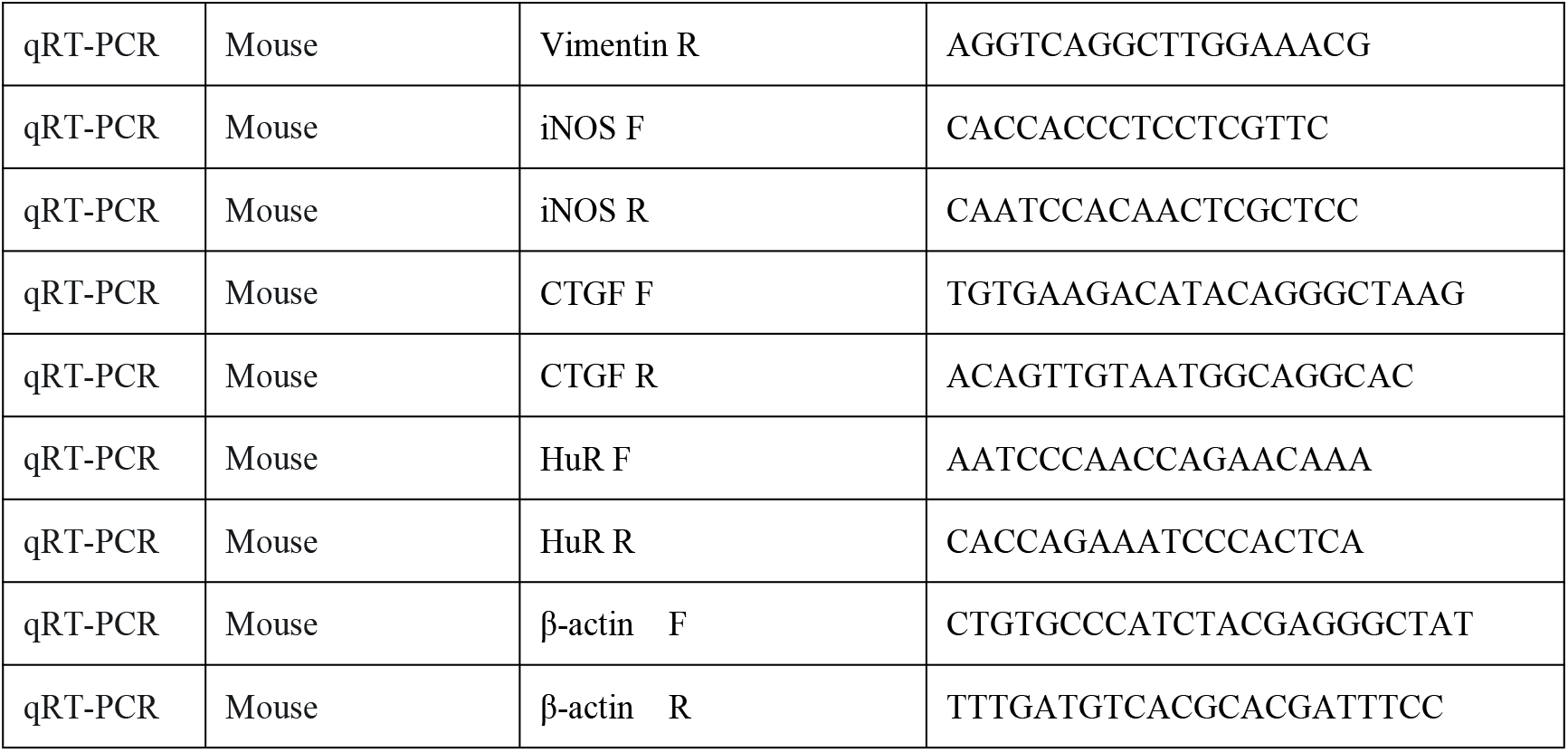
A list of the sequences of PCR primers.

**Supplemental method 3.** Primary antibodies and their working concentrations in Western blot (WB), immunofluorescence (IF) and immunohistochemistry (IHC).

| **Antibody** | **Producer** | **Identifier** | **Material** | **Usage** | **Concentrations** |
| --- | --- | --- | --- | --- | --- |
| Snail1 | Abclonal | A5243 | Lung tissue | WB | 1:500 |
| HuR | Proteintech | 11910-1-AP | Lung tissue | WB | 1:1000 |
| α-SMA | Abcam | ab7817 | Lung tissue | WB | 1:5000 |
| Collagen I | Abclonal | A21059 | Lung tissue | WB | 1:500 |
| Vimentin | Abclonal | A15596 | Lung tissue | WB | 1:500 |
| Twist1 | Abcam | ab50887 | Lung tissue | WB | 1:500 |
| HuR | Abcam | ab200342 | Lung tissue | IHC | 1:500 |
| α-SMA | Abcam | ab7817 | Lung tissue | IHC | 1:300 |
| HuR | Abcam | ab200342 | Lung tissue | IF | 1:200 |
| α-SMA | Abcam | ab7817 | Lung tissue | IF | 1:500 |
| osteopontin | Abcam | ab63856 | Lung tissue | IF | 1:100 |
| HuR | Abcam | ab200342 | Fibroblast | IF | 1:500 |
| α-SMA | Abcam | ab7817 | Fibroblast | IF | 1:300 |
| Snail1 | Abclonal | A5243 | Fibroblast | IF | 1:100 |
| osteopontin | Affinity | DF6395 | Fibroblast | IF | 1:100 |
| HuR | Proteintech | 11910-1-AP | Fibroblast | IP | 1:3000 |
| Fibronectin | Abcam | ab2413 | Fibroblast | WB | 1:500 |
| α-SMA | Abcam | ab7817 | Fibroblast | WB | 1:5000 |
| Collagen I | Abclonal | A21059 | Fibroblast | WB | 1:500 |
| Vimentin | Abclonal | A15596 | Fibroblast | WB | 1:500 |
| Snail1 | Abclonal | A5243 | Fibroblast | WB | 1:500 |
| p-Smad3 | Abclonal | AP0727 | Fibroblast | WB | 1:1000 |
| Smad3 | Abclonal | A19115 | Fibroblast | WB | 1:500 |
| osteopontin | Affinity | DF6395 | Fibroblast | WB | 1:1000 |
| HuR | Proteintech | 11910-1-AP | Fibroblast | WB | 1:1000 |

## Notes

### Competing Interest Statement

The authors have declared no competing interest.

